# The role of YAP/TAZ signaling in dendritic cell-mediated pathogenesis of insulin resistance and non-alcoholic fatty liver disease

**DOI:** 10.1101/2024.07.16.603813

**Authors:** Alejandro Schcolnik-Cabrera, Megan Lee, Kasia Dzierlega, Paulo Basso, Erin Strachan, Mengyi Zhu, Masoud Akbari, Xavier Clemente-Casares, Sue Tsai

## Abstract

Obesity and insulin resistance (IR) are global health challenges linked to metabolic diseases, such as type 2 diabetes and non-alcoholic fatty liver disease (NAFLD). High-caloric intake, which is associated to NAFLD, induces adipocyte hypertrophy and inflammation, triggering dendritic cell (DC) activation and systemic inflammation. DC exacerbate inflammation by promoting pro-inflammatory responses, aggravating IR and NAFLD progression. NAFLD is characterized by liver fibrosis, which alters tissue stiffness that can trigger mechanosensing pathways such as the Hippo pathway in immune cell types. In this work we explored the roles of key mediators of the Hippo pathway, YAP and TAZ, in DCs within the context of liver fibrosis, obesity and IR, using a model of NAFLD induced by feeding a high fat high sucrose diet. Our findings indicate that specific deletion of YAP and/or TAZ in DCs had minimal impact on IR development and metabolic tissue inflammation. We conclude that YAP and TAZ have limited and possibly redundant roles in the immune pathophysiology of NAFLD and IR.

## Introduction

Globally, obesity and insulin resistance (IR) pandemics affect over 650 million[1] and 1.24-3.7 billion subjects[2], respectively, predisposing them to chronic metabolic diseases like type 2 diabetes (T2D) and non-alcoholic fatty liver disease (NAFLD)[3,4]. Although the causes of the referred conditions are multidimensional in origin, key drivers include high visceral adiposity and associated long-term high-caloric intake (LTHCI)[5].

LTHCI induces adipocyte hypertrophy and inflammation via endoplasmic reticulum (ER) stress, releasing adipokines and chemoattractants that recruit and activate pro-inflammatory immune cells like macrophages and dendritic cells (DCs)[6]. Adipose tissue macrophages are polarized towards a pro-inflammatory phenotype, releasing cytokines that impair insulin signaling including TNF-α, IL-1β and IL-6 [7]. DCs worsen systemic inflammation by promoting Th1 and Th17 inflammatory responses[8], aggravating IR which, in turn, predisposes subjects to develop NAFLD[9].

NAFLD is a pro-inflammatory disorder characterized by liver fibrosis, which alters tissue stiffness that can potentially activate mechanosensing pathways such as the Hippo pathway[10,11]. Among other roles, the Hippo pathway has been shown to regulate cell survival, proliferation, migration and differentiation[12]. Yes-associated protein (YAP) and Transcriptional co-Activator with PDZ-binding motif (TAZ)[13] are key Hippo pathway transcriptional co-activators that are mobilized from the cytosol to the nucleus where they interact when unphosphorylated with the transcriptional enhanced associate domain (TEAD) transcription factors[14]. Particularly in DCs, YAP and TAZ sense extracellular mechanical and nutrient cues in DCs[15]. Under energy stress, AMPK phosphorylates both YAP and TAZ, blocking their activity. However, high glycemic levels inhibit AMPK, allowing YAP and TAZ to promote immune cell recruitment and activation[16].

Previous reports from us and others have demonstrated that substrate stiffness is an important determinant of DC function[17,18]. Indeed, in a high stiffness environment, DCs increase their activation and secretion of pro-inflammatory cytokines. Even though activated YAP/TAZ have been associated to the pathophysiology of highly inflammatory conditions such as liver cirrhosis and myocardial infarction, whether YAP/TAZ participate in DC mechanosensing and contribute to obesity-associated NAFLD is unknown.

Here, we investigated the role of YAP and TAZ signaling in DCs associated with liver fibrosis, obesity and IR. We generated DC-specific conditional YAP/TAZ gene deleted mice (CD11cCre^+^*Yap*^fl/fl^: YAP^DC-KO^; CD11cCre^+^*Wwtr1*^fl/fl^: TAZ^DC-KO^; and CD11cCre^+^*Yap*^fl/fl^: *Wwtr1*^fl/fl^: YAP/TAZ^DC-DKO^), and compared with their corresponding CD11cCre^−^ controls (CD11cCre^−^*Yap*^fl/fl^: YAP^DC-WT^; CD11cCre^−^ *Wwtr1*^fl/fl^: TAZ^DC-WT^; and CD11cCre^−^*Yap*^fl/fl^ *Wwtr1*^fl/fl^: YAP/TAZ^DC-WT^). While DC-specific ablation of YAP or TAZ did not significantly alter myeloid cell populations, YAP/TAZ^DC-DKO^ mice showed decreased macrophage and DC frequencies in the liver. However, YAP and/or TAZ single genetic ablation did not significantly impact IR development in high-fat high-sucrose diet (HFHS)-fed mice, and immune cell populations in livers and visceral adipose tissue (VAT) remained similar to their respective controls. Thus, YAP and TAZ mechanosensitive signaling plays limited and redundant roles in NAFLD pathophysiology.

## Results

### DC-specific YAP and TAZ deletion does not affect the myeloid cell population in the liver at steady-state

We first assessed the level of genetic deletion in respective YAP and TAZ KO mice. RT-qPCR analysis of isolated bone marrow dendritic cells (BMDCs) of these mice showed a 50% and 80% expression in YAP^DC-KO^ and YAP/TAZ^DC-DKO^, respectively, while *Wwtr1* expression was suppressed by 75% in both TAZ^DC-KO^ and YAP/TAZ^DC-DKO^ mice (**Supp. Fig. 1A-1D**).

We next profiled the myeloid cell population in livers at steady state in DC-specific YAP and/or TAZ KO mice. When comparing DC-specific deficiency of YAP, there were no differences in terms of frequency or absolute numbers between YAP^DC-KO^ and YAP^DC-WT^ mice, nor between TAZ^DC-KO^ and TAZ^DC-WT^ mice. In the case of the YAP/TAZ^DC-DKO^ mice, the proportion of liver monocytes, as well as the total numbers of monocyte-derived macrophages (MdM) and the conventional DCs type 1 and 2 (cDC1 and cDC2, respectively), were marginally decreased when compared against their YAP/TAZ^DC-WT^ control (**Fig. 1**, **Supp. Fig. 2**). These results suggest that YAP and TAZ deficiency may play a limited role in the development and/or migration of CD11c^+^ cells, comprising both DCs and MdMs cells, into the liver at steady state.

**Fig. 1:**
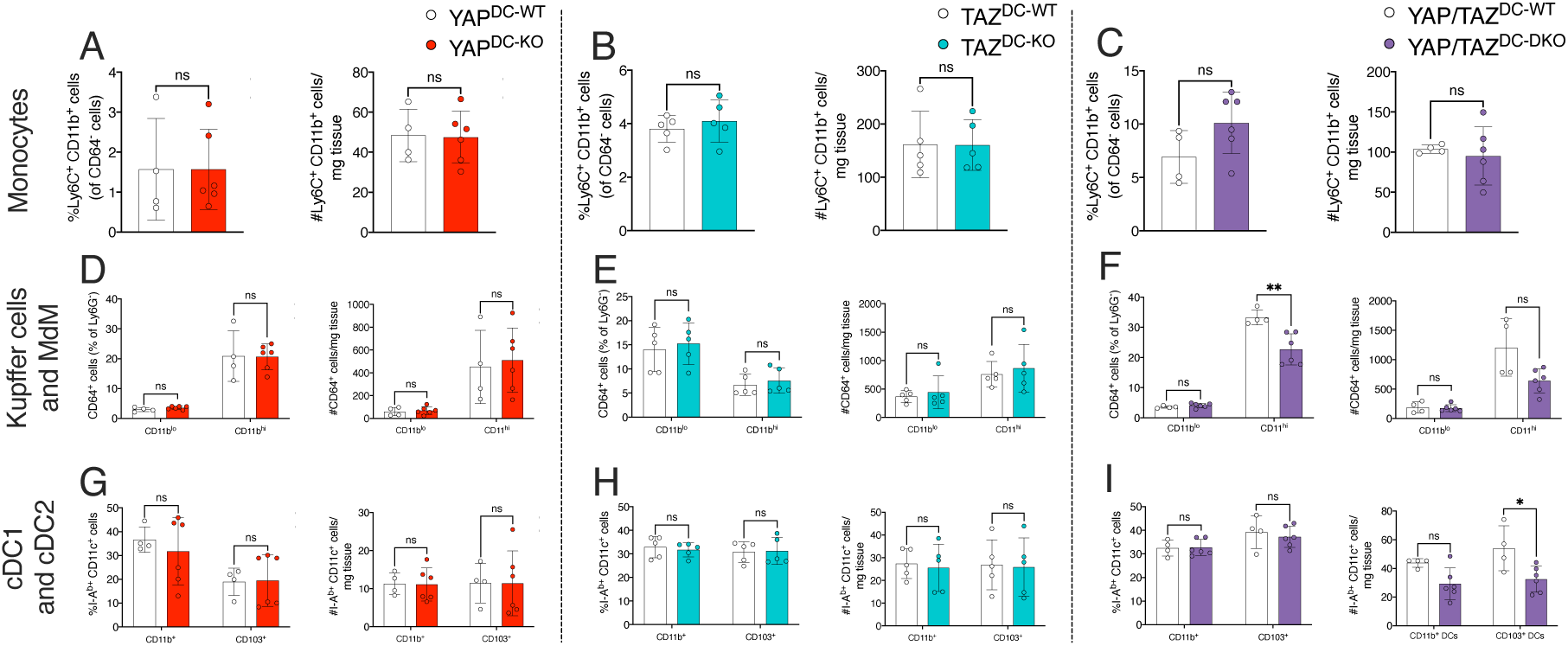
DC-specific YAP/TAZ deletion decreases the proportion of MdMs and cDC1s in mice liver at steady-state. Frequency (left) and absolute numbers (right) of monocytes in DC-specific YAP (**A**), TAZ (**B**) and YAP/TAZ (**C**) deletion. Frequency (left) and absolute numbers (right) of Kuppfer cells and MdMs in DC-specific YAP (**D**), TAZ (**E**) and YAP/TAZ (**F**) deletion. Frequency (left) and absolute numbers (right) of cDC1 and cDC2 in DC-specific YAP (**G**), TAZ (**H**) and YAP/TAZ (**I**) deletion. *Monocytes: Ly6C^+^CD11b^+^CD64^−^ cells; Kupffer cells: CD11b^lo^CD64^+^Ly6G^−^ cells; MdMs: CD11b^hi^CD64^+^Ly6G^−^ cells; cDC1: CD103^+^CD11c^+^CD64^−^I-A^b+^ cells; cDC2: CD11b^+^CD11c^+^CD64^−^I-A^b+^ cells. Number of cells was quantified and normalized to mg of tissue digested. Each dot represents an individual mouse. Data is shown as means±SD. Two-tailed Mann-Whitney U test. ns: non-significant; *p<0.05; **p<0.01*.

### DC YAP and/or TAZ deletion do not affect the development of insulin resistance in high fat/high sucrose fat-treated mice

Since obesity increases nutrient signals and alters tissue stiffness systemically including the liver[19], we hypothesized that DC YAP/TAZ signaling contributes to the obesity-IR axis. To address the hypothesis, we subjected YAP and TAZ single vs double KO mice to 20 weeks of diet regimen consisting of 60% kcal fat diet supplemented with 42 g/L of sucrose water (HFHS), starting at 6 weeks of age. Simultaneously, age-matched mice fed with a 4% kcal diet (normal chow diet; NCD) were employed as lean controls.

As expected, our data showed an increased in body weight gain in HFHS versus NCD-fed DC YAP/TAZ^WT^ mice (**Fig. 2A**). Furthermore, YAP and/or TAZ genetic ablation did not significantly alter body weight gain in mice fed a HFHS diet (**Fig. 2B-2C**). Next, to evaluate the glycemic responses to the administration of exogenous insulin and glucose, we performed insulin tolerance tests (ITT) and intraperitoneal glucose tolerance tests (IPGTT), respectively. The fasting blood glucose levels in NCD-fed mice were significantly lower compared to HFHS-fed mice across all the groups (**Suppl. Fig. 3A, 3C, 3E, Suppl. Fig. 3B, 3D, 3F**). When challenged with an intraperitoneal injection of insulin, NCD mice demonstrated significantly increased insulin sensitivity compared to HFHS-fed mice across all groups. However, no differences were observed between the YAP and TAZ DC KO groups (**Fig. 2D-2I**). Similarly, HFHS mice across all groups showed no difference in glucose tolerance (**Fig. 2J-2O**), other than a mildly increased fasting glucose in HFHS YAP^DC-KO^ vs YAP^DC-WT^ mice, and mildly improved glucose tolerance in HFHS TAZ^DC-KO^ vs TAZ^DC-WT^ mice. Overall, our results show that YAP and/or TAZ deletion in CD11c^+^ cells did not impact the development of obesity-associated IR in HFHS-fed mice.

**Fig. 2:**
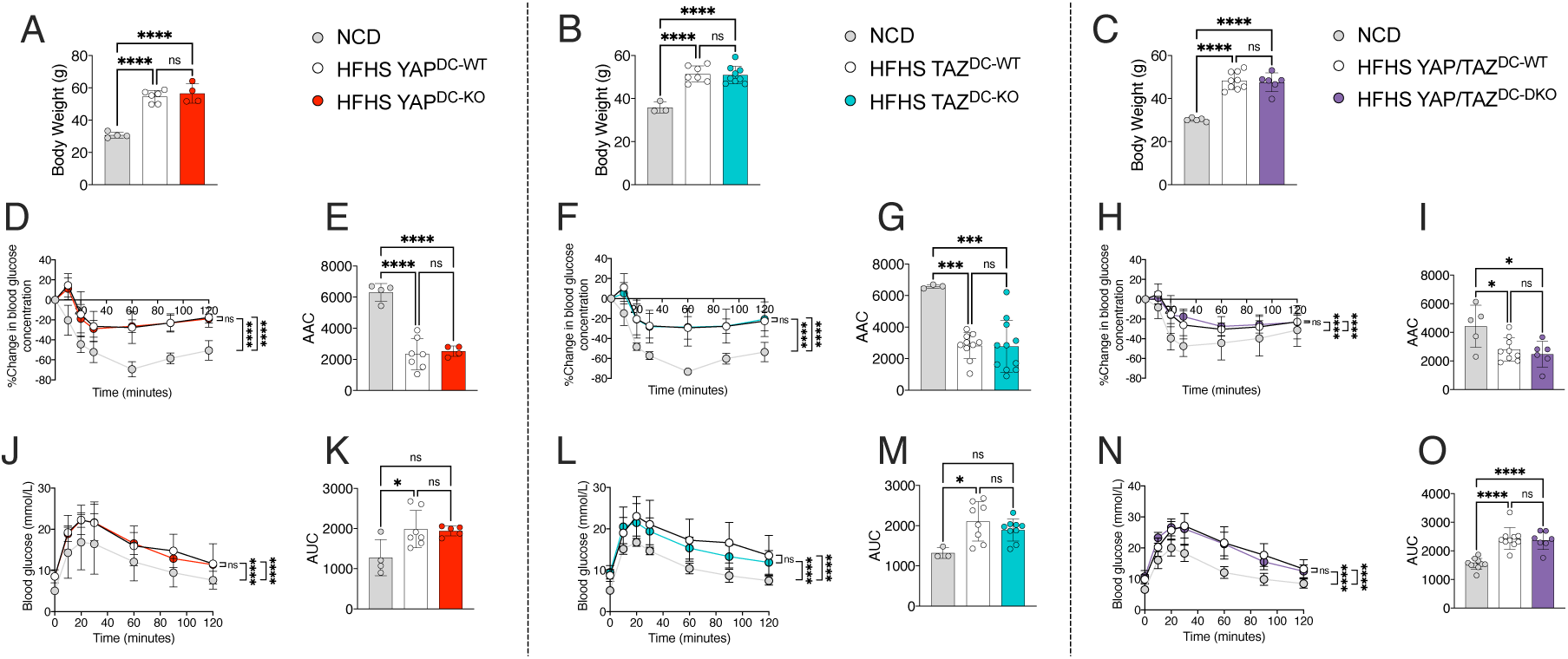
CD11c^+^-specific YAP and/or TAZ deletion do not affect the development of insulin resistance in HFHS-fed mice. Body weight measurement at *t*=20 weeks of feeding in DC-specific YAP (**A**), TAZ (**B**) or YAP/TAZ deletion (**C**) mice with either a NCD or a HFHS diet. Glucose excursion during ITT in DC-specific YAP (**D**), TAZ (**F**) or YAP/TAZ deletion (**H**) mice and their respective integrated area above the curve (AAC) (**E**, **G** and **I**, respectively). Glucose curve during IPGTT in DC-specific YAP (**J**), TAZ (**L**) or YAP/TAZ deletion (**N**) mice and their respective integrated area under the curve (AUC) (**K**, **M** and **O**, respectively). *For ITT, blood glucose is expressed as a percent change in glycemia from time point 0. Each dot represents an individual mouse. Data is shown as means±SD. Glucose excursion and glucose curve for ITT and IPGTT, respectively, were analyzed by two-way ANOVA with Tukey post-hoc test. The remaining analyses followed a one-way ANOVA with Tukey post-hoc test. ns: non-significant; *p<0.05; ***p<0.001; ****p<0.0001*.

### Deletion of YAP and/or TAZ in CD11c^+^ cells do not affect the levels of myeloid cells in livers of HFHS-treated mice

*YAP1* and *WWTR1* have been shown to be overexpressed across multiple liver pathologies in both human and mice, as demonstrated by a survey in the Gepliver database (http://www.gepliver.org) (**Supp. Fig 4A-4D**). Since YAP and TAZ signaling in DCs have been shown to boost their production of pro-inflammatory cytokines[17], we next aimed to investigate whether and how YAP and TAZ in DCs contribute to obesity-associated inflammation and the development of NAFLD.

After 20 weeks of HFHS diet feeding, mice exhibited liver inflammation when compared against NCD groups, as determined by an upregulation in the frequency and absolute numbers of immune cells in the liver. There was a marginal increase in the numbers of CD45^+^ cells and cDC2s in HFHS YAP^DC-WT^ and YAP^DC-KO^ mice (**Supp. Fig. 5A** and **Fig. 3G**), in the numbers of CD45^+^ cells and monocytes in HFHS TAZ^DC-WT^ and TAZ^DC-KO^ mice (**Supp. Fig. 5B** and **Fig. 3B**), and in the frequency of monocytes and in the MdM:Kupffer cells (KCs) ratio in HFHS YAP/TAZ^DC-WT^ and YAP/TAZ^DC-DKO^ mice (**Fig. 3C** and **Supp. Fig. 5I**). In addition, significant decreases were seen in frequencies and total numbers of neutrophils in HFHS YAP/TAZ^DC-WT^ and YAP/TAZ^DC-DKO^ mice (**Supp. Fig. 3F**). In both HFHS YAP^DC-WT^ and YAP^DC-KO^, and in HFHS YAP/TAZ^DC-WT^ and YAP/TAZ^DC-DKO^ mice, a trend to decrease in Kupffer cells was observed (**Fig. 3D, 3F**). However, when comparing HFHS-treated groups with CD11c-specific deletion of YAP and/or TAZ to their respective HFHS YAP and/or TAZ sufficient controls, our data did not show significant differences in the immune cell infiltration. Altogether, we report that YAP and/or TAZ deficiency in DCs does not significantly alter liver infiltration of myeloid cells during HFHS-induced obesity in mice.

**Fig. 3:**
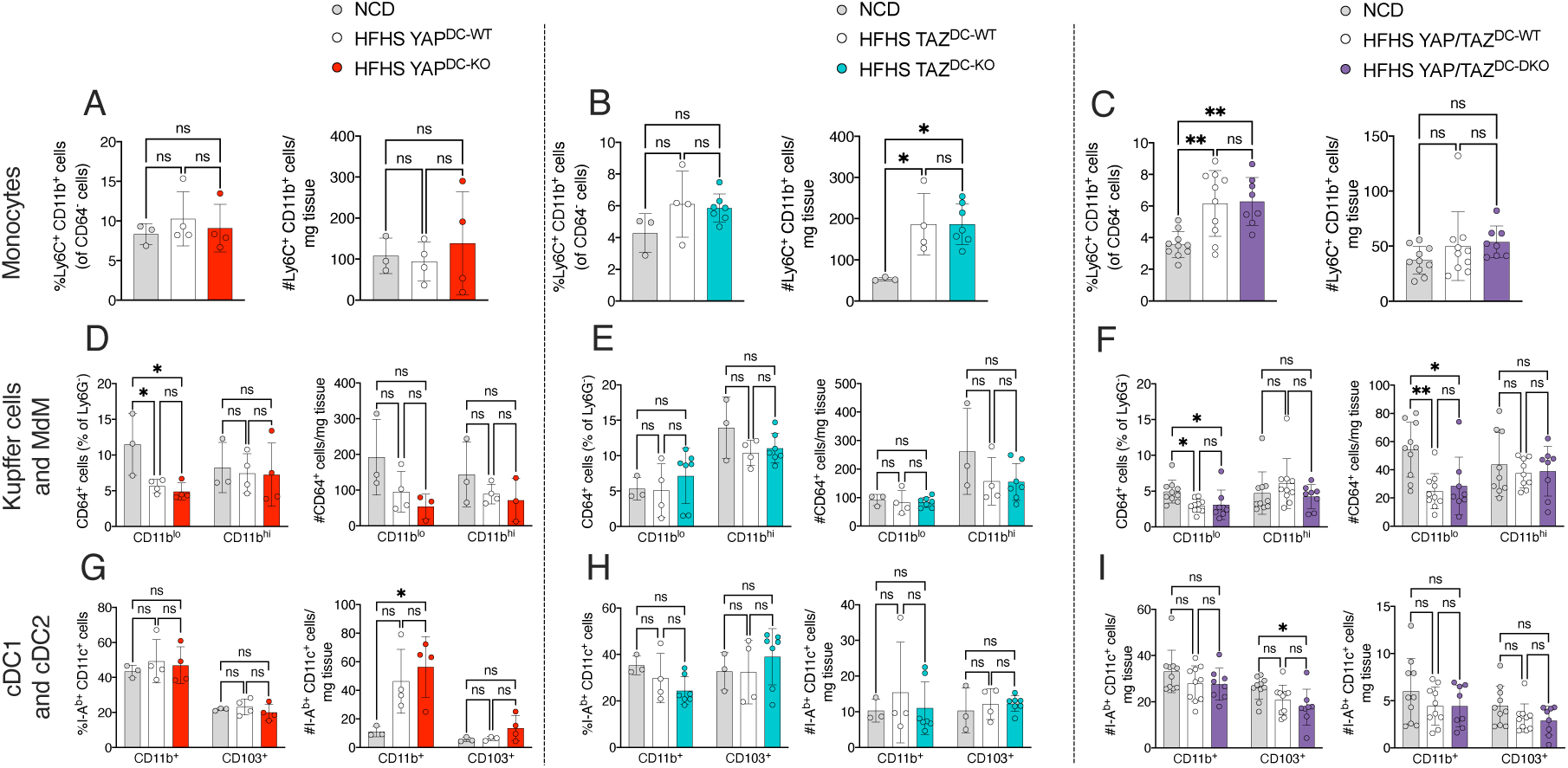
DC-specific YAP and/or TAZ deletion do not alter the levels of myeloid cell infiltration into the livers of HFHS-fed mice. Frequency (left) and absolute numbers (right) of monocytes in DC-specific YAP (**A**), TAZ (**B**) and YAP/TAZ (**C**) deletion. Frequency (left) and absolute numbers (right) of Kuppfer cells and MdMs in DC-specific YAP (**D**), TAZ (**E**) and YAP/TAZ (**F**) deletion. Frequency (left) and absolute numbers (right) of cDC1 and cDC2 in DC-specific YAP (**G**), TAZ (**H**) and YAP/TAZ (**I**) deletion. *Monocytes: Ly6C^+^CD11b^+^CD64^−^ cells; Kupffer cells: CD11b^lo^CD64^+^Ly6G^−^ cells; MdMs: CD11b^hi^CD64^+^Ly6G^−^ cells; cDC1: CD103^+^CD11c^+^CD64^−^I-A^b+^ cells; cDC2: CD11b^+^CD11c^+^CD64^−^I-A^b+^ cells. Number of cells was quantified and normalized to mg of tissue digested. Each dot represents an individual mouse. Data is shown as means±SD. One-way ANOVA with Tukey post-hoc test. ns: non-significant; *p<0.05; *p<0.05; **p<0.01*.

### T cell infiltration in the livers and VAT of HFHS-treated mice is unaffected by deletion of YAP and/or TAZ in CD11c^+^ cells

Obesity is accompanied by pro-inflammatory T cell infiltration in the liver and adipose tissues (AT) such as VAT[20,21], which is likely potentiated by DCs given their critical role in antigen presentation and cytokine secretion[22]. Given this premise, we next set out to evaluate the presence and activity of infiltrated T cells in both liver and VAT under YAP and/or TAZ deficiency in DCs of HFHS-fed mice.

Starting with the liver, our results an increase in the frequency and absolute numbers of total T cells in HFHS-fed mice vs NCD-fed mice (**Supp. Fig. 6A-B**). Both HFHS-fed YAP^DC-KO^ and HFHS-fed YAP^DC-WT^ mice showed altered CD4^+^ and CD8^+^T cell compartments, when compared against their NCD controls (**Supp. Fig. 6D, 6G**). Across the HFHS conditions, however, no differences in total T cells were seen. Flow cytometric analysis of the liver infiltrating immune cells revealed no significant changes in the populations of naïve (CD62L^hi^CD44^lo^), central memory (CD62L^hi^CD44^hi^), and effector memory T cells (CD44^hi^CD62L^lo^) between HFHS-treated mice with CD11c deletion of YAP and/or TAZ (**Fig. 4A-4F**). Thus, YAP and/or TAZ deficiency in DCs did not impact the T cell activation profiles in obese livers.

**Fig. 4:**
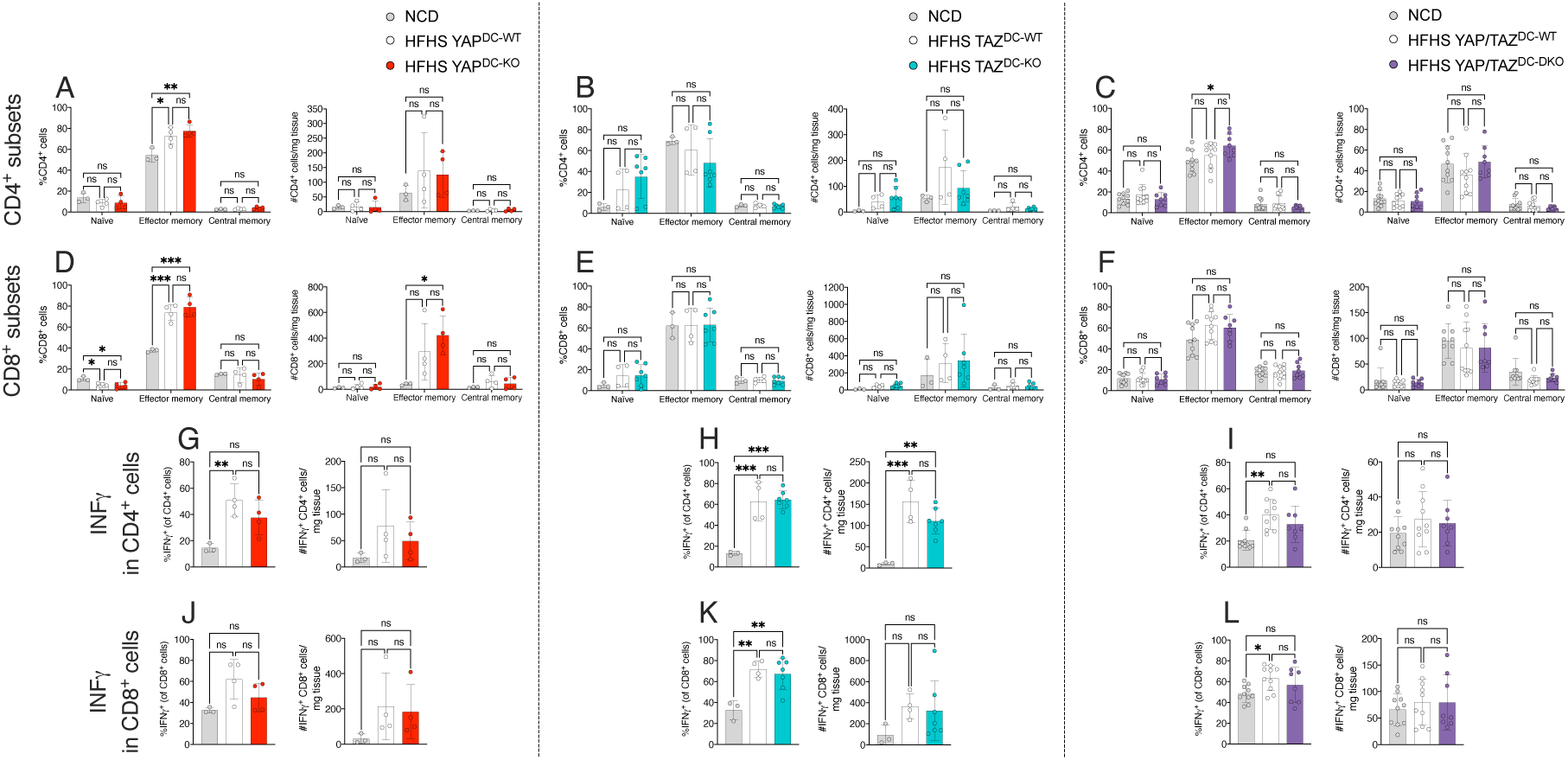
CD11c^+^-specific YAP and/or TAZ deletion do not modify the production of T cell pro-inflammatory cytokines in the livers of HFHS-fed mice. Frequency (left) and absolute numbers (right) of CD4^+^ T cell subsets in DC-specific YAP (**A**), TAZ (**B**) and YAP/TAZ (**C**) deletion. Frequency (left) and absolute numbers (right) of CD8^+^ T cell subsets in DC-specific YAP (**D**), TAZ (**E**) and YAP/TAZ (**F**) deletion. Frequency (left) and absolute numbers (right) of IFNγ^+^ CD4^+^ T cells in DC-specific YAP (**G**), TAZ (**H**) and YAP/TAZ (**I**) deletion. Frequency (left) and absolute numbers (right) of IFNγ^+^ CD8^+^ T cells in DC-specific YAP (**J**), TAZ (**K**) and YAP/TAZ (**L**) deletion. *Number of cells was quantified and normalized to mg of tissue digested. Each dot represents an individual mouse. Data is shown as means±SD. One-way ANOVA with Tukey post-hoc test. ns: non-significant; *p<0.05; *p<0.05; **p<0.01; ***p<0.001*.

In obese animals, liver CD8^+^ T cells produce increased amounts of TNFα- and IFNγ T cells driven by cDC1s, and IFNγ- and IL17-secreting CD4^+^T cells, stimulated by cDC2s [23,24]. After stimulating both liver-derived CD4^+^ and CD8^+^ populations with PMA and ionomycin, the livers of HFHS-fed mice showed a trending upregulation in the proportion and numbers of IFNγ-producing CD8^+^T, and in IFNγ- and IL17-secreting CD4^+^T cells, but only when compared against NCD-fed mice (**Fig. 4G-4L** and **Supp. Fig. 6J-6L**). Similarly, a mild and non-significant increase in TNFα-producing CD8^+^T cells was seen in HFHS-fed YAP^DC-KO^, HFHS-fed YAP^DC-WT^, HFHS-fed YAP/TAZ^DC-DKO^, and HFHS-fed YAP/TAZ^DC-WT^ mice, when compared against NCD mice. Following the same trend as with the total T cell populations, no differences in the frequencies and numbers of IFNγ^+^CD4^+^T cells, IL17^+^CD4^+^T cells, or IFNγ^+^CD8^+^T cells, were seen in HFHS-fed mice associated with CD11c ablation of YAP and/or TAZ. However, only HFHS YAP^DC-KO^ mice demonstrated a tendency towards reduced frequency of IFNγ-producing CD4^+^ and CD8^+^T cells against their HFHS YAP^DC-WT^ control mice (**Fig. 4G, 4J**). This suggests that in DCs, YAP activity moderately modulates Th1 differentiation.

Lastly, the composition of VAT-infiltrated immune cells was also assessed in HFHS-fed mice, as VAT is a critical site of inflammation related to obesity. CD11c-specific deletion of YAP and/or TAZ was not associated with altered frequencies of CD45^+^ cells, neutrophils (**Supp. Fig. 7A-7B, 7D-7E, 7G-7H**) and monocytes (**Fig. 5A, 5D, 5G**). Similarly, comparable proportions of CD11c^−^CD206^+^ and CD11c^+^CD206^+^ macrophages, comprising anti-inflammatory and pro-inflammatory AT-infiltrated macrophages[25,26], were seen in CD11c-specific deletion of YAP and/or TAZ mice (**Fig. 5J, 5L, 5N**). No differences were seen regarding the proportions of cDC1s and cDC2s (**Fig. 5K, 5M, 5O**), nor the CD4^+^T or CD8^+^T cell compartments (**Fig. 5B-5C, 5E-5F, 5H-5I**), or their populations of naïve, effector memory, and central memory (**Fig. 5P-5U**), in the VAT from with CD11c-specific deletion of YAP and/or TAZ fed with a HFHS diet. Only a minor and not significant decrease in the frequency of total T cells in HFHS YAP/TAZ^DC-DKO^ and HFHS TAZ^DC-KO^ mice (**Supp. Fig. 7F, 7I**), and in the proportion of CD8^+^ effector memory T cells in HFHS TAZ^DC-KO^ mice (**Fig. 5S**), was seen against their controls. Altogether, our data show that YAP and/or TAZ deletion in DCs does not alter immune cell infiltration in the VAT during obesity.

**Fig. 5:**
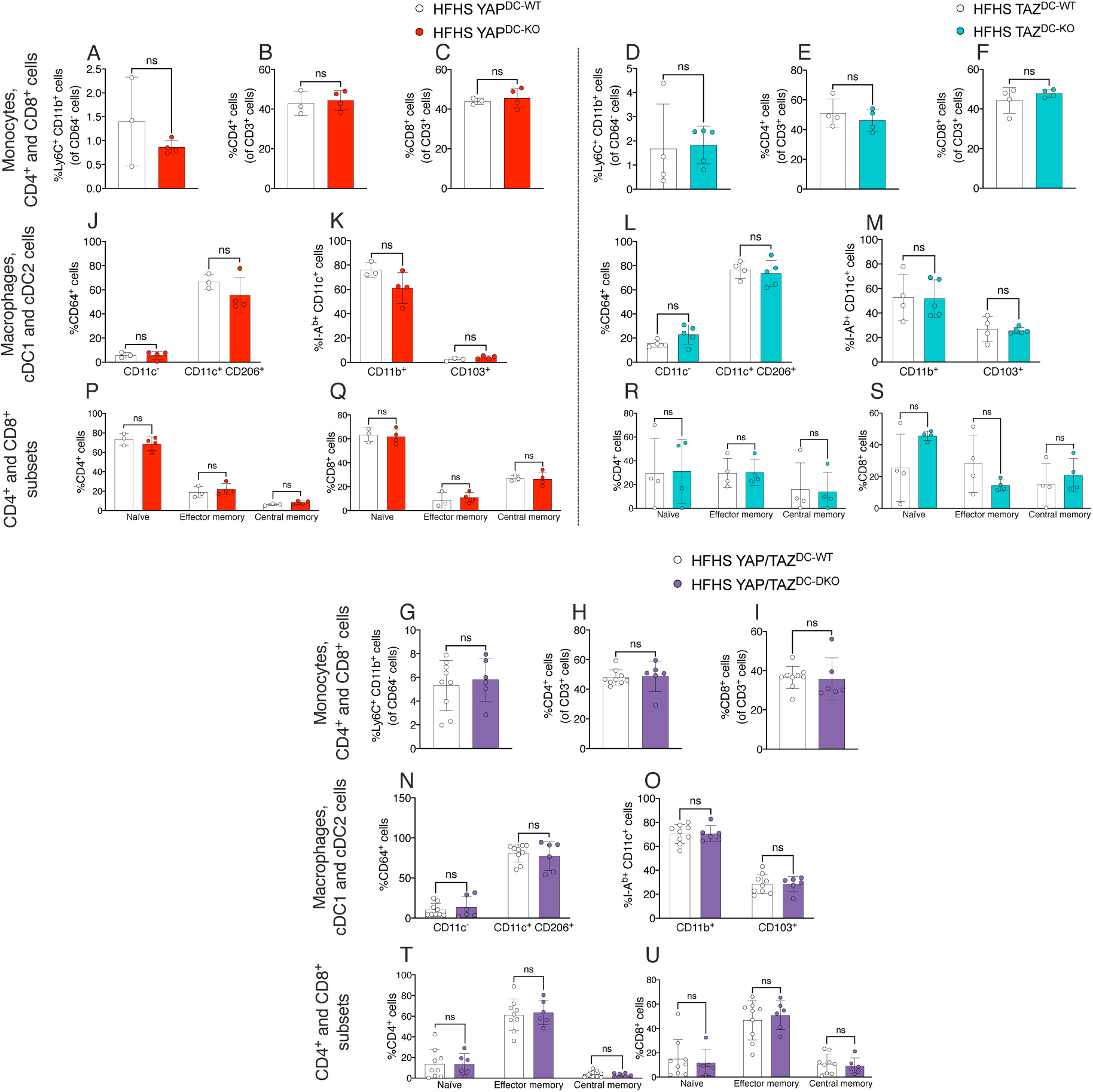
DC-specific YAP and/or TAZ deletion do not affect the frequencies of myeloid and T cell population in the VATs of HFHS-fed mice. Frequencies of monocytes, CD4^+^ T and CD8^+^ T cells in DC-specific YAP (A-C), TAZ (D-F) and YAP/TAZ (G-I) deletion. Frequencies of macrophages, cDC1 and cDC2 in DC-specific YAP (J-K), TAZ (L-M) and YAP/TAZ (N-O) deletion. Frequencies of CD4^+^ T (left) and CD8^+^ T (right) cell subsets in DC-specific YAP (P-Q), TAZ (R-S) and YAP/TAZ (T-U) deletion. *Monocytes: Ly6C^+^CD11b^+^CD64^−^ cells; anti-inflammatory VAT-resident macrophages: CD11c^−^CD64^+^ cells; pro-inflammatory, IR-associated, inflammation-related VAT resident macrophages: CD11c^+^CD206^+^CD64^+^ cells; cDC1: CD103^+^CD11c^+^CD64^−^I-A^b+^ cells; cDC2: CD11b^+^CD11c^+^CD64^−^I-A^b+^ cells. Each dot represents an individual mouse. Data is shown as means±SD. Two-tailed Mann-Whitney U test. ns: non-significant*.

### Genetic deletion of YAP and/or TAZ in CD11c^+^ cells did not ameliorate liver fibrosis in HFHS-fed mice

Besides progressive liver inflammation, a critical determinant of NAFLD is liver fibrosis associated to extracellular collagen deposition, which occurs as a wound-healing mechanism against repeated injury[27]. When comparing our HFHS-fed YAP/TAZ^DC-DKO^ and HFHS-fed YAP/TAZ^DC-WT^ mice, similar proportions of collagen fibers were stained by Sirius red in liver histology samples (**Supp. Fig. 7J-7L**). These results suggest that the combined YAP and TAZ deletion in DCs does not influence the natural progression of fibrosis in the context of HFHS and NAFLD.

## Discussion

Using a CD11cCre model, we investigated the roles of YAP and TAZ signaling in DCs associated to NAFLD in HFHS-fed mice. Despite the known roles of YAP and TAZ in cell cycle regulation and proliferation[28], our observations revealed no significant differences in the hepatic myeloid cell population between YAP^DC-KO^ and TAZ^DC-KO^ mice and their control littermates under steady-state conditions. However, in the double knockout model, we observed a reduction in the proportions and numbers of MdMs and cDC1s cells in livers.

While previous studies have shown that YAP expression increases during monocyte differentiation into macrophages[29], investigations using *Yap*^fl/fl^/*Taz*^fl/fl^ mice indicated no alterations in the numbers of circulating white blood cells over time or in the efficiency of bone marrow cell differentiation into myeloid and lymphoid lineages[30]. However, considering that DCs precursors migrate from the bone marrow to peripheral tissues for terminal differentiation into cDCs, it is conceivable that the effects observed in the liver under the double knockout condition may be attributed to the roles of YAP/TAZ in DC development, migration, and liver homing. Although not specific to DCs, Fan S. et al. demonstrated in multiple myeloma cells that overexpression of YAP or TAZ did not affect cell proliferation but promoted transwell migration, a phenomenon associated with suppression of mitophagy[31]. Additionally, the authors reported enrichment of TAZ mRNA in bone marrow-derived CD138^+^ cells from multiple myeloma patients, with higher expression levels of YAP or TAZ correlating with reduced patient survival [31]. Thus, further investigation is warranted to elucidate the roles of YAP/TAZ in DC migration and tissue homing, particularly in the liver environment.

It has been indicated that YAP/TAZ signaling in DCs is associated with increased production of pro-inflammatory cytokines[17]. Given the link between obesity-associated inflammation and IR development, we hypothesized that YAP and TAZ signaling in DCs might impact IR development. Compared to their NCD control counterparts, HFHS-fed YAP^DC-KO^, TAZ^DC-KO^, and YAP/TAZ^DC-DKO^ mice demonstrated significantly higher fasting glycemic levels, suggesting IR development related to diet. However, similar insulin sensitivity and glucose tolerance were observed across CD11c-specific deletion of YAP and/or TAZ mice compared to their HFHS-fed CD11cCre^−^ controls, suggesting that DC YAP and/or TAZ deletion does not affect IR development. The observed phenotypes may be associated with additional mechanisms involved in obesity-associated IR, such as increased circulating levels of free fatty acids (FFAs), which activate JNK and IKK50, leading to phosphorylation of serine residues on the insulin receptor substrate (IRS) and its subsequent inactivation. Additional players in the inactivation cascade include pro-inflammatory macrophages, which increase in number and activity during obesity, further stimulating JNK and IKK via TNF-α secretion, inhibiting IRS[32]. Furthermore, FFAs taken up by adipose tissue-resident DCs have a dual effect on promoting antigen presentation. They activate the MAPK pathway, inducing MHC-II expression[33], and they enhance CD4^+^ and CD8^+^T cell activity through lipid droplet-induced antigen cross-presentation[34]. However, whether high circulating levels of FFAs directly impact DC activity in the context of YAP and/or TAZ deletion, requires further evaluation.

While VAT inflammation is a key driver for systemic inflammation in obesity, liver inflammation indicates progression to NAFLD[35]. In our study, immune cell infiltration levels in both VAT and livers of HFHS-fed YAP^DC-KO^, TAZ^DC-KO^, and YAP/TAZ^DC-DKO^ mice were comparable to their respective controls. Interestingly, livers of HFHS-fed YAP^DC-KO^ mice showed a trending decrease in proportions of IFNγ^+^CD4^+^T and IFNγ^+^CD8^+^T cells compared to HFHS-fed YAP^DC-WT^ control mice. These results suggest that YAP signaling in DCs may regulate IL-12 and IFNγ production, which are necessary for Th1 differentiation[36]. To explore this hypothesis, further functional comparison of cytokine production by YAP-deficient DCs against YAP-sufficient DCs is essential.

Lastly, we assessed liver fibrosis levels in HFHS-fed YAP/TAZ^DC-DKO^ and YAP/TAZ^DC-WT^ mice, as a measure of NAFLD progression. The Fibrosis-4 Index (FIB) is an accepted approach to evaluate advanced liver fibrosis[37]. In NAFLD patients, there is a direct correlation between increasing liver stiffness levels and the FIB-4 score, demonstrating progressive histological severity of fibrosis with liver stiffness >10kPa[38]. Certainly, ≥20% rise in liver stiffness, or ≥5kPa increase, acts as a predictive factor in patients for NALFD progression into cirrhosis[39]. In our study, we found similar levels of tissue fibrosis, consistent with analogous levels of inflammation and inflammatory cell populations in the livers of both groups. While these findings suggest that YAP and/or TAZ signaling in DCs may not significantly contribute to NAFLD pathogenesis, evaluating additional NAFLD markers such as homocysteine and glutathione levels, as well as the AST/platelet ratio and FIB-4 score index[40], would enhance our understanding of this phenotype.

The role of YAP/TAZ in integrating various extracellular signals beyond mechanical stiffness are well-established. YAP/TAZ are known to respond to a range of microenvironmental cues, including nutrient availability and inflammatory signals[41]. In endothelial cells, YAP and/or TAZ regulate angiogenesis and amino acid uptake through mTORC1 signaling, which depends on nutrient levels [42]. Additionally, in environments characterized by high adhesion and stiffness that promote cytoskeletal polymerization, YAP expression can drive macrophage inflammation [29]. Given that the degree of liver tissue stiffness in our models relative to controls is currently undetermined, it is plausible that YAP/TAZ signaling may be only partially activated due to inadequate changes in local tissue tissues. Consequently, the cumulative effects of YAP/TAZ in modulating extracellular cues might obscure the impact of stiffness in our experimental model. Further experiments are necessary to elucidate the precise relationship between tissue stiffness in NAFLD, and DC-specific YAP and/or TAZ deletion.

## Abbreviations

IR: insulin resistance
T2D: type 2 diabetes
NAFLD: non-alcoholic fatty liver disease
LTHCI: long-term high-caloric intake
ER: endoplasmic reticulum
DCs: dendritic cells
YAP: Yes-associated protein
TAZ: Transcriptional co-Activator with PDZ-binding motif
TEAD: transcriptional enhanced associate domain
YAP^DC-KO^: CD11cCre^+^*Yap*^fl/fl^ mice
TAZ^DC-KO^: CD11cCre^+^*Yaz*^fl/fl^ mice
YAP/TAZ^DC-DKO^: CD11cCre^+^*Yap*^fl/fl^*Taz*^fl/fl^ mice
YAP^DC-WT^: CD11cCre^−^*Yap*^fl/fl^ mice
TAZ^DC-WT^: CD11cCre^−^*Yaz*^fl/fl^ mice
YAP/TAZ^DC-WT^: CD11cCre^−^ *Yap*^fl/fl^*Taz*^fl/fl^ mice
HFHS: high-fat high-sucrose diet
VAT: visceral adipose tissue
MdM: monocyte-derived macrophages
cDC1: conventional dendritic cells type 1
cDC2: conventional dendritic cells type 2
NCD: normal chow diet
ITT: insulin tolerance test
IPGTT: intraperitoneal glucose test
KCs: Kupffer cells
FFAs: free fatty acids
IRS: insulin receptor substrate
PMNs: polymorphonuclear cells
AAC: area above the curve
AUC: area under the curve

## Acknowledgments

ASC would like to thank the NSERC-CREATE program for the postdoctoral fellowship granted.

## Author contributions

S.T.: original conceptualization; A.S.-C., M.L., K.D., P.B., E.S., M.Z., M.A.: experimentation, mice handling, data analysis; A.S.-C.: figure preparation; A.S.-C.: first manuscript draft; A.S.-C., S.T.: revision of the manuscript; S.T., X.C.-C.: supervision of the study; S.T.: resources provision and funds acquisition. All authors have approved the final version of the manuscript.

## Materials and methods

### Mice

All mice were generated by intercrossing breeders obtained from Jackson Mice. Genetically manipulated mice were generated as described in Supp. Fig. 1A. 8-12 week old male and female mice were used. All mice were bred and housed in a pathogen-free, temperature-controlled, and 12h light/dark cycle environment at the University of Alberta Health Sciences Laboratory Animal Services mouse barrier facility. All mice were gender- and age-matched. Mice were fed *ad libitum* with either a normal chow diet (NCD, 4% fat) or with a HFHS diet, as indicated next. All experimental procedures were approved by the Animal Care committee at the University of Alberta. Mice genotypes were confirmed by PCR and gel electrophoresis.

### Diet-induced obesity (DIO) model

6 week-old male C57BL/6 mice were fed a HFD consisting of 60% fat for 16 weeks[43]. IPGTT and ITT were tested following 16 weeks of feeding to assess for glucose tolerance and insulin tolerance, respectively.

### HFHS model

6 week-old mice were put on a HFD consisting of 60% fat, supplemented with 42g/L sucrose water for 20 weeks. IPGTT and ITT were conducted following 20 weeks of feeding to assess for glucose tolerance and insulin tolerance, respectively. Male mice were used for HFHS studies as male mice are more susceptible to diet-induced weight gain and IR[44].

### BDMC culture

BMDCs were cultured following Inaba *et al*.[45]. Briefly, bone marrow progenitor cells were harvested from the femur of C57BL/6 mice and plated at a density of 3×10^6^/3mL well on 6-well plates with 20 ng/mL GM-CSF (BioLegend) in RPMI 1640 medium (Wisent) supplemented with 10% fetal bovine serum (FBS) (Sigma), 2mM L-glutamine (Gibco), 100 U/mL penicillin (Sigma), 100 μg/mL streptomycin (Sigma), 50μM 2-mercaptoethanol (Sigma), 1mM sodium pyruvate (Gibco), and MEM non-essential amino acids (Gibco). Bone marrow cultures were fed on days 3 and 6 with 1mL cell culture medium containing fresh GM-CSF.

### Real-time PCR

Total RNA from cultured BMDCs was extracted using the PureLink^TM^ RNA mini kit (ThermoFisher) as per manufacturer’s protocol. Reverse-transcription of the RNA was performed using random primers (ThermoFisher). qPCR was performed on a CFX96 Touch Real-Time PCR Detection System (BioRad) using SUPERGREEN^TM^ Master Mix reagent (Wisent). Each sample was run in triplicate and normalized to housekeeping gene β−actin. Relative fold changes in gene expression were calculated using the ΔΔCT method with the equation 2^−ΔΔCT^. The results are shown as fold changes compared to the control group average. For genetic floxing efficiency of *Yap1* and *Wwtr1* (WW domain containing transcription regulator 1; also known as Taz), primers specific for detecting exon 2 of *Yap1* and *Wwtr1* were used. Sequences of the mouse forward and reverse primers are as follows: *β−actin* forward: 5’-GACCTCTATGCCAACACAGT-3’; reverse: 5’-AGTACTTGCGCTCAGGAGGA-3’; *Yap1* forward: 5’-TACTGATGCAGGTACTGCGG-3’; reverse: 5’-TCAGGGATCTCAAAGGAGGAC-3’; *Wwtr1* forward: 5’-CAAGTCATCCACGTCACGCA-3’; reverse: 5’-CCGGAATCGGGCTCCTTAAA-3’. The complete list of commercial assay and kits used for this study can be found in **Supplementary Table 1**.

### IPGTT and ITT

#### IPGTT

Mice were fasted overnight for 16h. Body weights were measured to calculate the dosage required to inject 1.5g glucose/kg body weight. In a biosafety cabinet (BSC) at the animal facility, mice were anesthetized under inhalation of 2% isoflurane. Next, the tail vein was gently punctured with a 25 gauge needle and a baseline blood glucose reading was acquired using a glucometer. Mice were intraperitoneally injected with glucose solution, and blood glucose excursion was measured at 10, 20, 30, 60, 90, and 120 min post-injection.

#### ITT

Mice were fasted for 6h. In a BSC at the animal facility, mice were anesthetized under inhalation of 2% isoflurane. Next, the tail vein was gently punctured with a 25 gauge needle and a baseline blood glucose reading was acquired using a glucometer. Mice were intraperitoneally injected with insulin (0.75 U insulin/kg body weight; Humalog), and blood glucose levels were measured at 10, 20, 30, 60, 90, and 120 minutes post-injection.

### Tissue isolation and digestion

Mice were euthanized by CO_2_ fixation, following the protocols provided by the University of Alberta Health Sciences Laboratory Animal Services. Liver tissue was minced and subjected to collagenase I digestion (450 U/mL; Sigma) with DNase-I (60 U/mL; Sigma) in 800μL of DMEM (Wisent) with gentle shaking for 1h at 37 °C, and filtered through a 40μm cell strainer with a HBB (HBSS, 2% bovine serum, 0.2% BSA) wash. After centrifugation (1500RPM, 5min) cell pellets were resuspended in 1mL of 40% Percoll® for density separation. Supernatant was aspirated and single cell suspensions were resuspended in ACK lysing buffer for 5 min. Epididymal VAT pads were dissected from mice and were mechanically dissociated using a gentleMACS Octo Dissociator in 5mL of DMEM, and then subjected to collagenase I digestion (2mg/mL; Sigma) with DNase-I (60 U/mL; Sigma) with gentle shaking for 1h at 37 °C, and filtered through a 40μm cell strainer with a PBS wash. The complete list of reagents used for this study can be found in **Supplementary Table 2**.

### Flow cytometry

Immune cells from mouse tissues were stained with a live/dead exclusion dye (Zombie UV, BioLegend). Fluorophore-conjugated antibodies were diluted 1:200 unless otherwise recommended by the supplier prior to surface and intracellular staining at 4 °C for 30min. Intracellular staining was done after fixation and permeabilization with a FOXP3 Staining Buffer Set, following manufacturer’s instructions (eBioscience). For intracellular cytokine staining, cell suspensions were stimulated with phorbol myristate acetate (PMA) and ionomycin in the presence of Brefeldin A (PMA+, BioLegend) for 5h. Antigen-specific CD4^+^T and CD8^+^T cells were stained with tetramers (NIH) specific for OVA_329-337_ H-2IA^b^ (AAHAEINEA). Permeabilized/fixed samples were resuspended in permeabilization buffer, while unfixed samples were resuspended in FACS buffer for flow cytometric analysis on a BD LSR Fortessa-SORP or Cytek Aurora at the Flow Cytometry Facility of the University of Alberta. FCS plots were generated using FlowJo software V10.10.0. The complete list of antibodies used for this study can be found in **Supplementary Table 3**.

### Liver fibrosis staining

One liver lobe was isolated from HFHS-fed mice, embedded in paraffin blocks, and longitudinally serially sectioned at a thickness of 7μm. Deparaffinization started with 100% xylene, followed by decrease ethanol concentrations until finally ddH_2_O. Then, tissues were stained with Sirius Red/Fast Green Collagen Staining Kit (Catalog #9046, Chondrex Inc) for 45min to identify collagen fibers suggestive of fibrosis development. Tissues were washed with ddH_2_O and rehydrated (from ddH_2_O to ethanol until 100% xylene), and mounted with a toluene mounting medium. Liver fibrosis was quantified as the proportion of liver positive for Sirius Red staining. Images were concluded using a Cytation C10 confocal microscope (Agilent), and staining intensity was quantified by ImageJ V1.51 (ImageJ, NIH).

### Statistical analysis

Unless otherwise indicated, all the reported experiments were performed in at least three independent assays. Statistical analysis were executed as indicated in each figure legend. All the results were analyzed and represented using GraphPad Prism V9.01 (GraphPad, San Diego, CA, USA). *p*<0.05 was considered statistically significant.

## Supplementary figure legends

**Supp. Fig. 1: Validation of *Yap1* and *Wwtr1* genetic ablation by RT-PCR in BMDCs in YAP^DC-KO^, TAZ^DC-KO^, and YAP/TAZ^DC-DKO^ mice**. Gene expression of *Yap1* (**A**), *Wwtr1* (**B**) and both *Yap1* and *Wwtr1* (**C**) from cultured BMDCs isolated from YAP^DC-KO^, TAZ^DC-KO^, and YAP/TAZ^DC-DKO^ mice, respectively. *Each dot represents an individual mouse. Data is shown as means±SD. Two-tailed Mann-Whitney U test. ns: non-significant; *p<0.05*.

**Supp. Fig. 2: DC-specific YAP/TAZ deletion decreases the MdMs/Kupffer cell ratio in mice liver at steady-state**. Frequency (left) and absolute numbers (right) of hematopoietic-derived cells in DC-specific YAP (**A**), TAZ (**B**) and YAP/TAZ (**C**) deletion. Frequency (left) and absolute numbers (right) of PMNs in DC-specific YAP (**D**), TAZ (**E**) and YAP/TAZ (**F**) deletion. MdMs/Kupffer cell ratio in DC-specific YAP (**G**), TAZ (**H**) and YAP/TAZ (**I**) deletion. *Hematopoietic-derived cells: CD45^+^ cells; PMNs: Ly6G^+^CD45^+^ cells; MdMs: CD11b^hi^CD64^+^Ly6G^−^ cells; Kupffer cells: CD11b^lo^CD64^+^Ly6G^−^ cells. Number of cells was quantified and normalized to mg of tissue digested. Each dot represents an individual mouse. Data is shown as means±SD. Two-tailed Mann-Whitney U test. ns: non-significant; *p<0.05*.

**Supp. Fig. 3: CD11c^+^-specific YAP and/or TAZ deletion do not modify fasting glycemia in HFHS-fed mice**. Fasting glycemia measurements at *t*=20 weeks of feeding in DC-specific YAP, TAZ, or YAP/TAZ deletion mice with either a NCD or a HFHS diet. Measurements at time point 0 after 6h of fasting for ITT and 16h of fasting for IPGTT in CD11c^+^-specific YAP (**A**, **B**), TAZ (**C**, **D**), or YAP/TAZ (**E**, **F**) deletion. *Each dot represents an individual mouse. Data is shown as means±SD. One-way ANOVA with Tukey post-hoc test. ns: non-significant; *p<0.05; **p<0.01; ***p<0.001; ****p<0.0001*.

**Supp. Fig. 4: *YAP1* and *WWTR1* levels are enhanced in human and mice liver diseases**. *YAP1* and *WWTR1* expression in livers from human (**A** and **C**, respectively) and mice (**B** and **D**, respectively) origin across liver pathologies. *Data was curated from the Gepliver database (*http://www.gepliver.org*). NAFLD: non-alcoholic fatty liver disease; ALD: alcohol-related liver disease; hepatocellular carcinoma; ICC: intrahepatic cholangiocarcinoma; HB: hepatitis B; IPNB: intraductal papillary neoplasm of the bile duct. Data distribution is shown as violin plots. One-way ANOVA with Tukey post-hoc test. **p<0.01; ***p<0.001; ****p<0.0001*.

**Supp. Fig. 5: DC-specific YAP and/or TAZ deletion do not modify the infiltration of hematopoietic cells including neutrophils and macrophages into the livers of HFHS-fed mice**. Frequency (left) and absolute numbers (right) of CD45^+^ cells in DC-specific YAP (**A**), TAZ (**B**) and YAP/TAZ (**C**) deletion. Frequency (left) and absolute numbers (right) of PMNs in DC-specific YAP (**D**), TAZ (**E**) and YAP/TAZ (**F**) deletion. MdMs/Kupffer cell ratio in DC-specific YAP (**G**), TAZ (**H**) and YAP/TAZ (**I**) deletion. *Hematopoietic-derived cells: CD45^+^ cells; PMNs: Ly6G^+^CD45^+^ cells; MdMs: CD11b^hi^CD64^+^Ly6G^−^ cells; Kupffer cells: CD11b^lo^CD64^+^Ly6G^−^ cells. Number of cells was quantified and normalized to mg of tissue digested. Each dot represents an individual mouse. Data is shown as means±SD. One-way ANOVA with Tukey post-hoc test. ns: non-significant; *p<0.05; **p<0.01; ***p<0.001*.

**Supp. Fig. 6: DC-specific YAP and/or TAZ deletion do not affect the frequencies of T cell population in the livers of HFHS-fed mice**. Frequency (left) and absolute numbers (right) of CD3^+^, CD4^+^, and CD8^+^ cells in DC-specific YAP (**A, D, G**), TAZ (**B, E, H**) and YAP/TAZ (**C, F, I**) deletion. Frequency (left) and absolute numbers (right) of IL-17 CD4^+^ cells in DC-specific YAP (**J**), TAZ (**K**) and YAP/TAZ (**L**) deletion. Frequency (left) and absolute numbers (right) of TNF-α CD8^+^ cells in DC-specific YAP (**M**), TAZ (**N**) and YAP/TAZ (**O**) deletion. *Number of cells was quantified and normalized to mg of tissue digested. Each dot represents an individual mouse. Data is shown as means±SD. One-way ANOVA with Tukey post-hoc test. ns: non-significant; *p<0.05; **p<0.01; ***p<0.001*.

**Supp. Fig. 7: DC-specific YAP and/or TAZ deletion do not influence hematopoietic cells infiltration in the VAT including neutrophils and T cells, nor induce liver fibrosis in HFHS-fed mice**. Frequencies of CD45^+^ cells, neutrophils and CD3^+^ cells in DC-specific YAP (**A-C**), TAZ (**D-F**) and YAP/TAZ (**G-I**) deletion isolated from the VAT. Liver section samples stained with Sirius Red/Fast Green from DC-specific YAP/TAZ^DC-WT^ (**J**) or YAP/TAZ^DC-DKO^ (**K**) mice at *t*=20 weeks of fed with HFHS diet. Liver fibrosis in both tested groups (**L**). *Hematopoietic-derived cells: CD45^+^ cells; PMNs: Ly6G^+^CD45^+^ cells. The histological images are representative from each tested group (bar=200μm). Each dot represents an individual mouse. Data is shown as means±SD. Two-tailed Mann-Whitney U test. ns: non-significant*.

## Supplementary Tables

**Supplementary Table 1.** List of commercial assays and kits.

| **Commercial assays/kits** | **Source** | **Catalog number** |
| --- | --- | --- |
| FOXP3/Transcription Factor Staining Buffer Set | eBioscience | 00-5523-00 |
| PureLink™ RNA mini kit | ThermoFisher | 12183018A |
| Random primers | ThermoFisher | 48190011 |
| Advanced qPCR Mastermix with SUPERGREEN™ | Wisent | 801-001-AR |
| Sirius red/Fast green Collagen staining kit | Chondrex, Inc. | 9046 |

**Supplementary Table 2.**
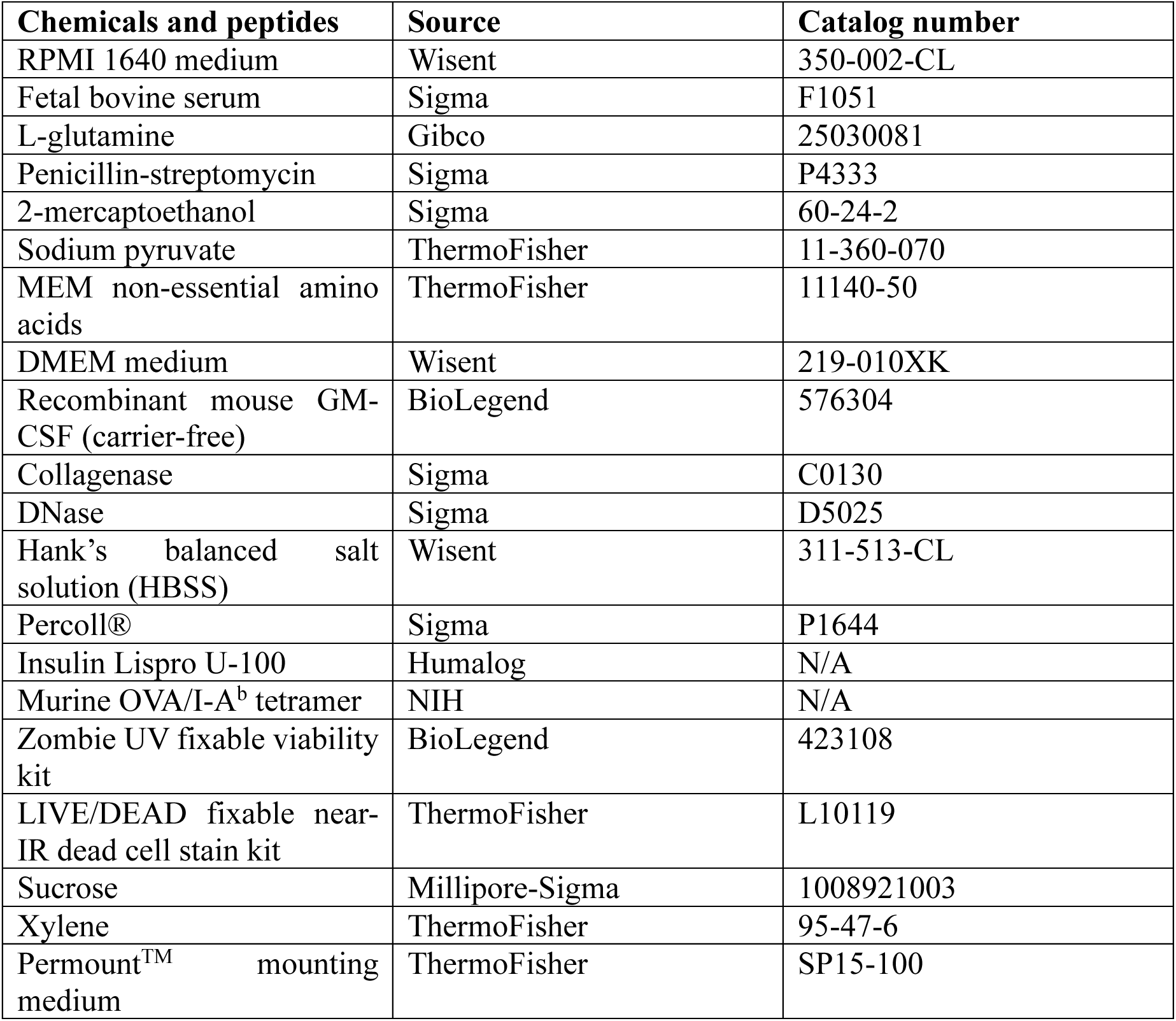
List of reagents.

**Supplementary Table 3.** List of antibodies for flow cytometry.

| **Antibody** | **Clone** | **Source** | **Catalog number** |
| --- | --- | --- | --- |
| Brilliant Violet 711 anti-mouse CD45 | 30-F11 | BioLegend | 103147 |
| Alexa Fluor 700 anti-mouse CD45 | I3/2.3 | BioLegend | 147716 |
| Alexa Fluor 488 anti-mouse CD3 | 17A2 | BioLegend | 100210 |
| PE/Dazzle 594 anti-mouse CD4 | GK1.5 | BioLegend | 100456 |
| PE/Cyanine5 anti-mouse CD8a | 53-6.7 | BioLegend | 100710 |
| Brilliant Violet 605 anti-mouse CD8a | 53-6.7 | BioLegend | 100744 |
| Brilliant Violet 510 anti-mouse CD62L | MEL-14 | BioLegend | 104441 |
| PerCP-Cy 5.5 anti-mouse FOXP3 | R16-715 | BD Biosciences | 563902 |
| PE anti-mouse IFNγ | XMG1.2 | BioLegend | 505808 |
| Alexa Fluor 700 anti-mouse Ly-6G | 1A8 | BioLegend | 127622 |
| APC/Cyanine7 anti-mouse Ly-6C | HK1.4 | BioLegend | 128026 |
| Brilliant Violet 605 anti-mouse CD11c | N418 | BioLegend | 117334 |
| FITC anti-mouse I-A<sup>b</sup> | AF6-120.1 | BioLegend | 116406 |
| APC anti-mouse CD103 | 2E7 | BioLegend | 121414 |
| PerCP/Cyanine5.5 anti-mouse/human CD11b | M1/70 | BioLegend | 101228 |
| PE/Cyanine7 anti-mouse CD64 (FcγRI) | X54-5/7.1 | BioLegend | 139314 |

