## Supplementary material for "The role of YAP/TAZ signaling in dendritic cell-mediated pathogenesis of insulin resistance and non-alcoholic fatty liver disease": Supplementary files_The role of YAP:TAZ signaling in dendritic cell-mediated pathogenesis of insulin resistance and non-alcoholic fatty liver disease.pdf

**Supp. Fig. 1**

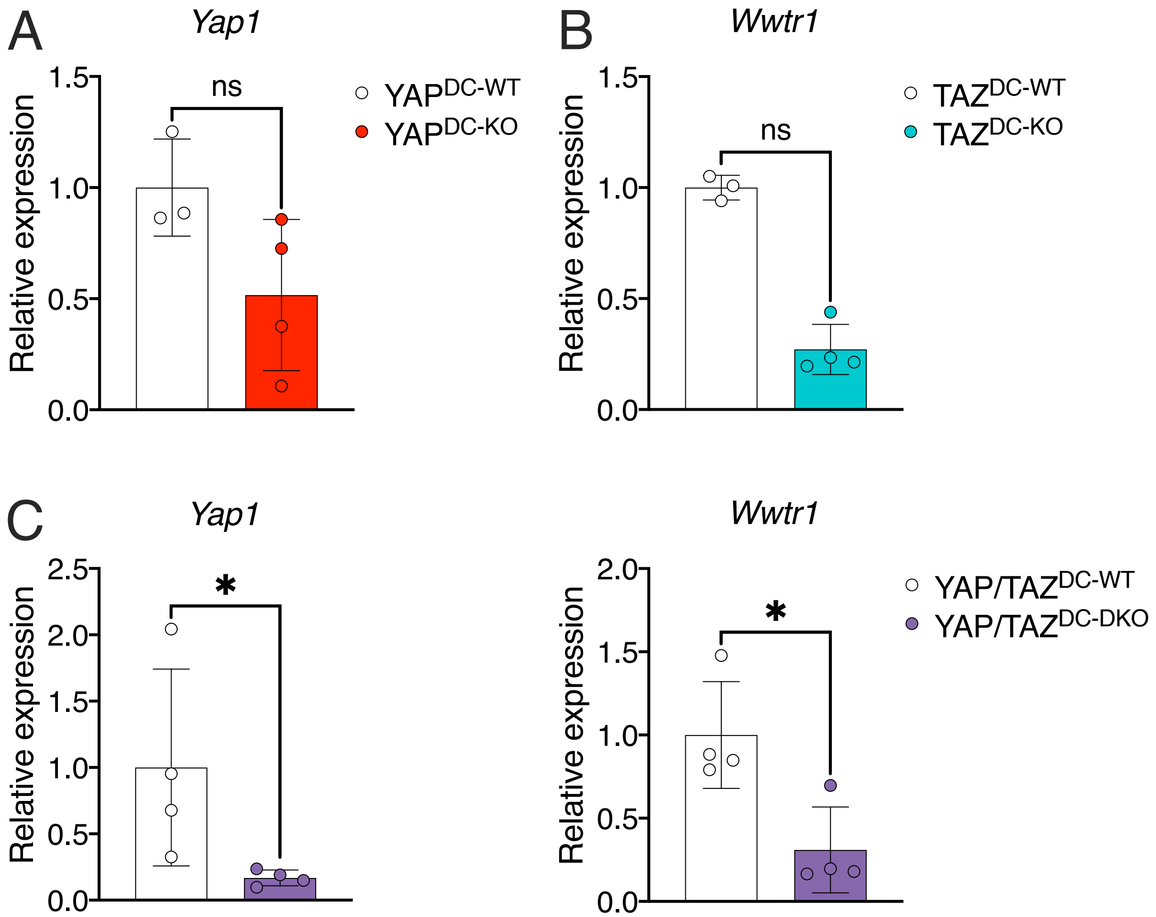
A
Yap1
Relative expression
1.5
1.0
0.5
0.0
ns
YAP^DC-WT
YAP^DC-KO
B
Wwtr1
Relative expression
1.5
1.0
0.5
0.0
ns
TAZ^DC-WT
TAZ^DC-KO
C
Yap1
Relative expression
2.5
2.0
1.5
1.0
0.5
0.0
*
Wwtr1
Relative expression
2.0
1.5
1.0
0.5
0.0
*
YAP/TAZ^DC-WT
YAP/TAZ^DC-DKO

**Supp. Fig. 2**

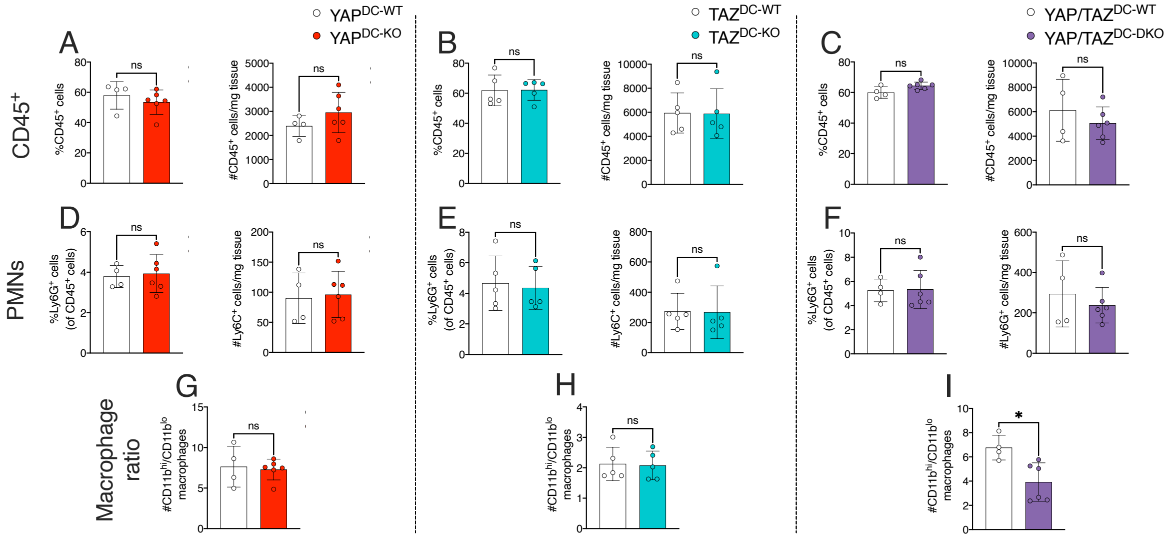

YAP^DC-WT
YAP^DC-KO
TAZ^DC-WT
TAZ^DC-KO
YAP/TAZ^DC-WT
YAP/TAZ^DC-DKO
A
B
C
D
E
F
G
H
I
CD45+
PMNs
Macrophage ratio
%CD45+ cells
#CD45+ cells/mg tissue
%Ly6G+ cells (of CD45+ cells)
#Ly6C+ cells/mg tissue
#Ly6G+ cells/mg tissue
#CD11b^hi/CD11b^lo macrophages
ns
*

**Supp. Fig. 3**

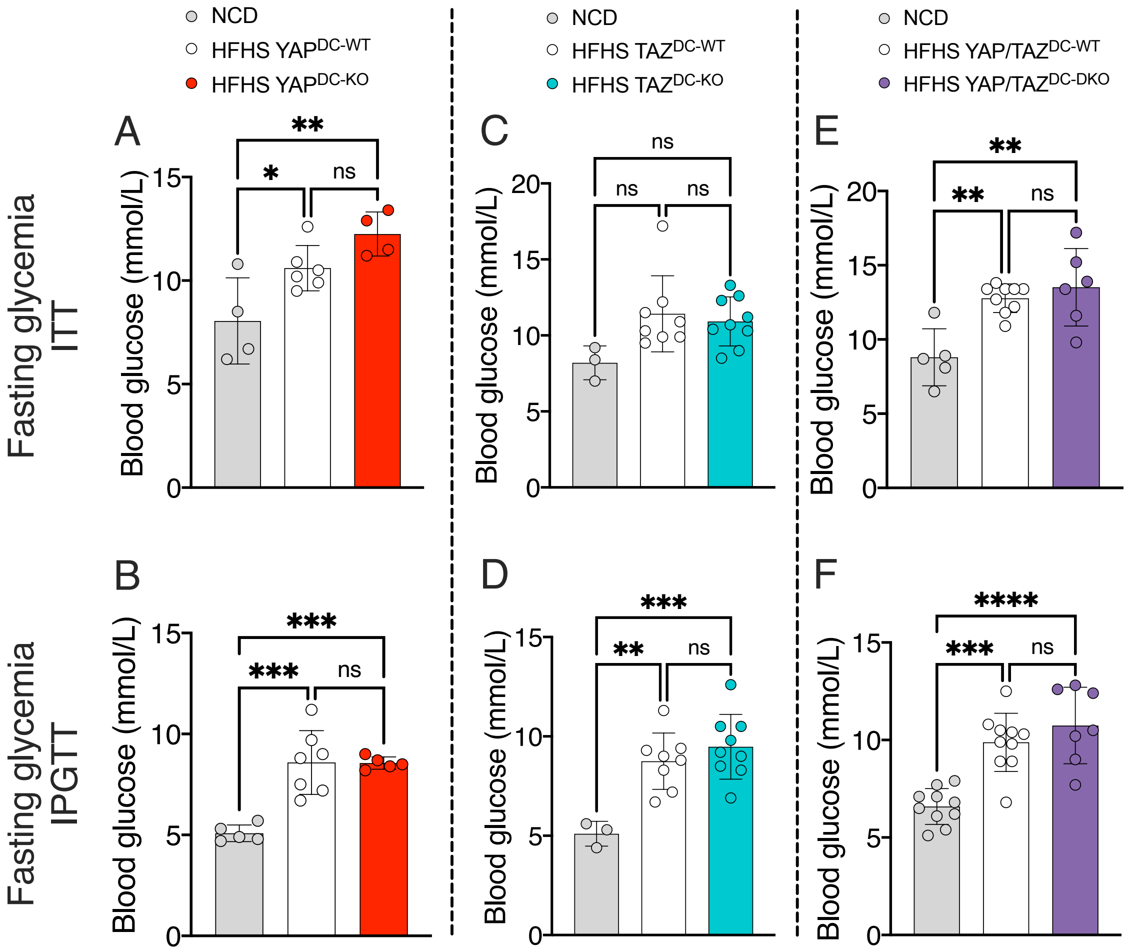

NCD
HFHS YAP$^{DC\text{-}WT}$
HFHS YAP$^{DC\text{-}KO}$
NCD
HFHS TAZ$^{DC\text{-}WT}$
HFHS TAZ$^{DC\text{-}KO}$
NCD
HFHS YAP/TAZ$^{DC\text{-}WT}$
HFHS YAP/TAZ$^{DC\text{-}DKO}$
Fasting glycemia ITT
Fasting glycemia IPGTT
A
B
C
D
E
F
Blood glucose (mmol/L)
ns

**Supp. Fig. 4**

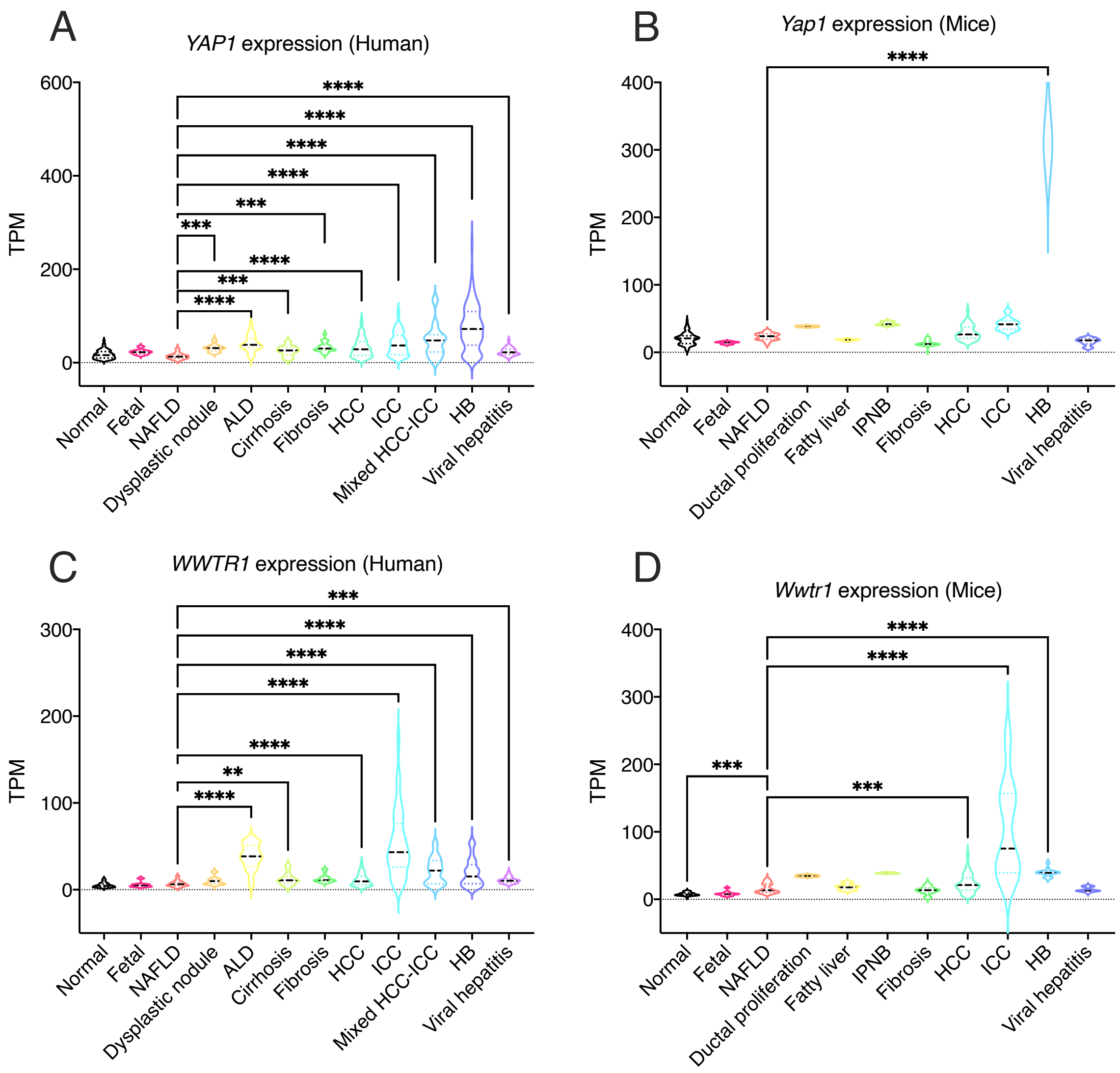
A
YAP1 expression (Human)
TPM
600
400
200
0
****
****
****
****
***
***
****
***
****
Normal
Fetal
NAFLD
Dysplastic nodule
ALD
Cirrhosis
Fibrosis
HCC
ICC
Mixed HCC-ICC
HB
Viral hepatitis
B
Yap1 expression (Mice)
TPM
400
300
200
100
0
****
Normal
Fetal
NAFLD
Ductal proliferation
Fatty liver
IPNB
Fibrosis
HCC
ICC
HB
Viral hepatitis
C
WWTR1 expression (Human)
TPM
300
200
100
0
***
****
****
****
****
**
****
Normal
Fetal
NAFLD
Dysplastic nodule
ALD
Cirrhosis
Fibrosis
HCC
ICC
Mixed HCC-ICC
HB
Viral hepatitis
D
Wwtr1 expression (Mice)
TPM
400
300
200
100
0
****
****
***
***
Normal
Fetal
NAFLD
Ductal proliferation
Fatty liver
IPNB
Fibrosis
HCC
ICC
HB
Viral hepatitis

**Supp. Fig. 5**

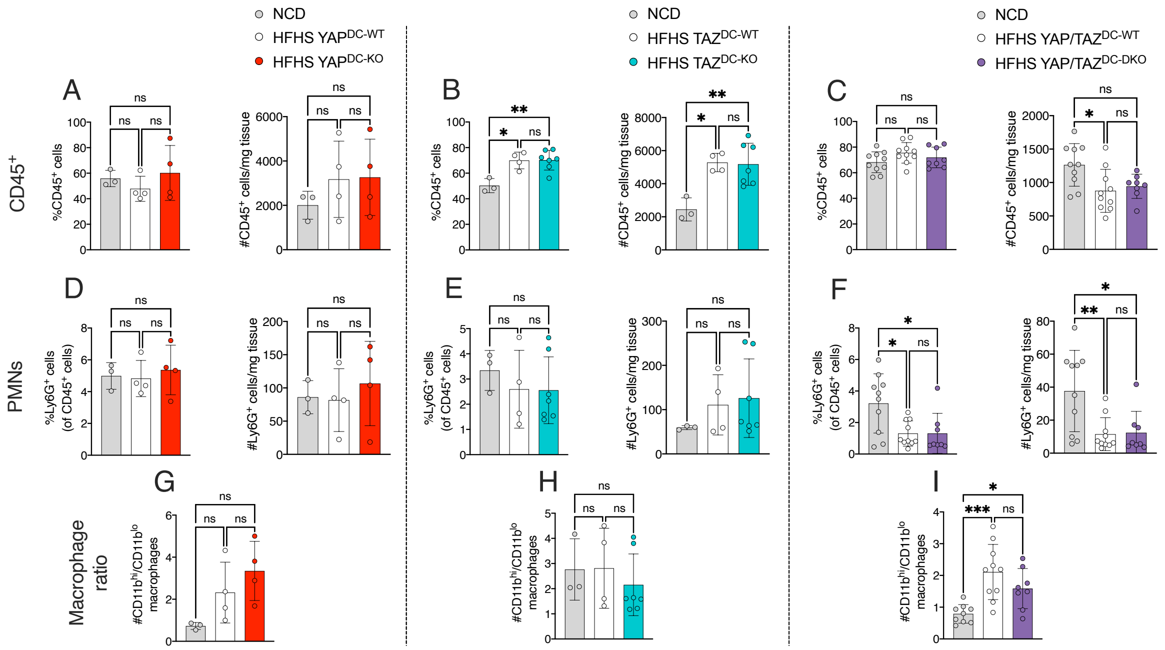

NCD
HFHS YAP$^{DC-WT}$
HFHS YAP$^{DC-KO}$
NCD
HFHS TAZ$^{DC-WT}$
HFHS TAZ$^{DC-KO}$
NCD
HFHS YAP/TAZ$^{DC-WT}$
HFHS YAP/TAZ$^{DC-DKO}$
A
B
C
D
E
F
G
H
I
CD45$^{+}$
PMNs
Macrophage ratio
%CD45$^{+}$ cells
#CD45$^{+}$ cells/mg tissue
%Ly6G$^{+}$ cells (of CD45$^{+}$ cells)
#Ly6G$^{+}$ cells/mg tissue
#CD11b$^{hi}$/CD11b$^{lo}$ macrophages

**Supp. Fig. 6**

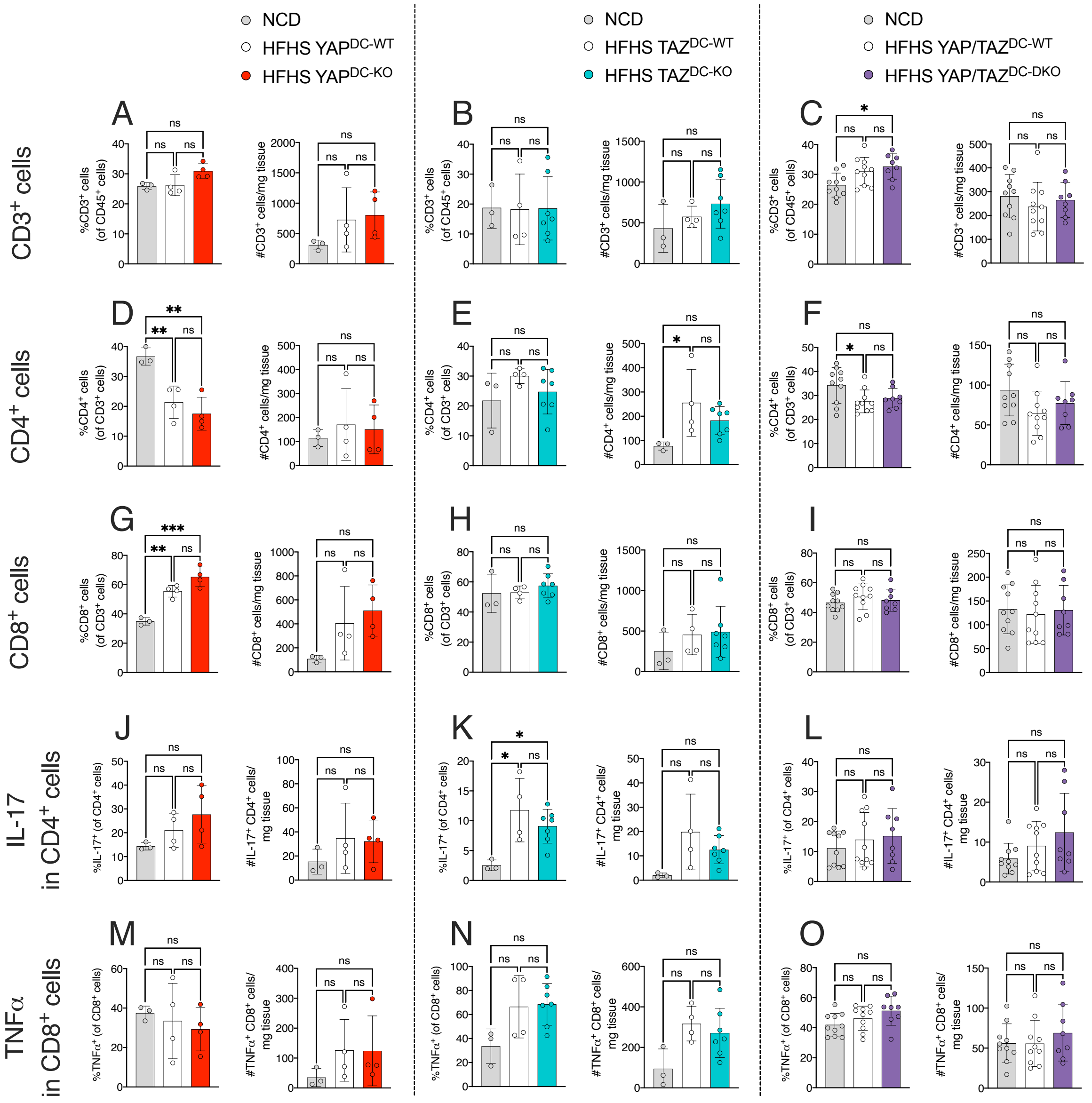
NCD
HFHS YAP$^{DC-WT}$
HFHS YAP$^{DC-KO}$
NCD
HFHS TAZ$^{DC-WT}$
HFHS TAZ$^{DC-KO}$
NCD
HFHS YAP/TAZ$^{DC-WT}$
HFHS YAP/TAZ$^{DC-DKO}$
CD3$^{+}$ cells
CD4$^{+}$ cells
CD8$^{+}$ cells
IL-17 in CD4$^{+}$ cells
TNFα in CD8$^{+}$ cells
A
B
C
D
E
F
G
H
I
J
K
L
M
N
O
%CD3$^{+}$ cells (of CD45$^{+}$ cells)
#CD3$^{+}$ cells/mg tissue
%CD4$^{+}$ cells (of CD3$^{+}$ cells)
#CD4$^{+}$ cells/mg tissue
%CD8$^{+}$ cells (of CD3$^{+}$ cells)
#CD8$^{+}$ cells/mg tissue
%IL-17$^{+}$ (of CD4$^{+}$ cells)
#IL-17$^{+}$ CD4$^{+}$ cells/ mg tissue
%TNFα$^{+}$ (of CD8$^{+}$ cells)
#TNFα$^{+}$ CD8$^{+}$ cells/ mg tissue

**Supp. Fig. 7**

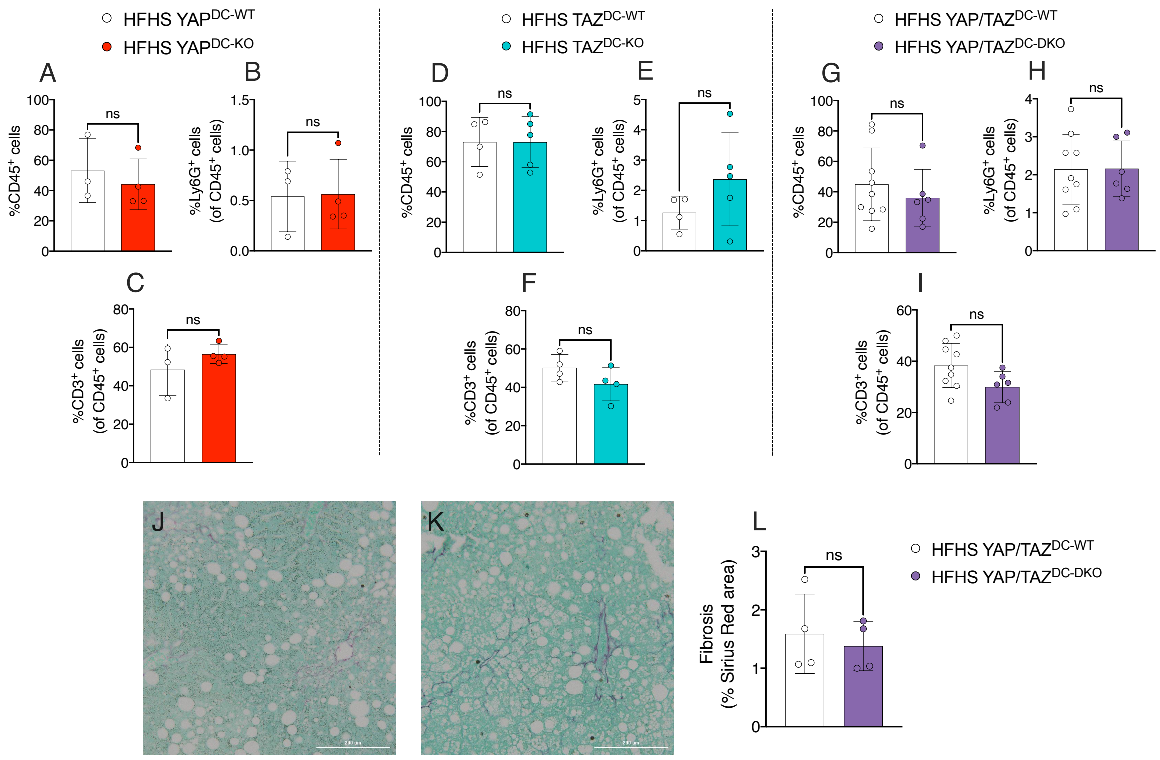
HFHS YAP^DC-WT
HFHS YAP^DC-KO
HFHS TAZ^DC-WT
HFHS TAZ^DC-KO
HFHS YAP/TAZ^DC-WT
HFHS YAP/TAZ^DC-DKO
A
B
C
D
E
F
G
H
I
J
K
L
ns
%CD45+ cells
%Ly6G+ cells (of CD45+ cells)
%CD3+ cells (of CD45+ cells)
Fibrosis (% Sirius Red area)
